# Mutation of the ALS/FTD-associated RNA-binding protein FUS alters axonal cytoskeletal organisation

**DOI:** 10.1101/2022.10.04.510780

**Authors:** Francesca W. van Tartwijk, Lucia C.S. Wunderlich, Ioanna Mela, Stanislaw Makarchuk, Maximilian A.H Jakobs, Seema Qamar, Kristian Franze, Gabriele S. Kaminski Schierle, Peter H. St George-Hyslop, Julie Qiaojin Lin, Christine E. Holt, Clemens F. Kaminski

**Author notes:** Friedrich-Alexander University Erlangen-Nuremberg, Institute of Medical Physics, Henkestr. 91, 91052 Erlangen, Germany; Max-Planck-Zentrum für Physik und Medizin, 91054 Erlangen, Germany (Kristian Franze).

## Abstract

Aberrant condensation and localisation of the RNA-binding protein fused in sarcoma (FUS) occur in variants of amyotrophic lateral sclerosis (ALS) and frontotemporal dementia (FTD). ALS is also associated with cytoskeletal defects, genetically and through observations of compromised axonal transport. Here, we asked whether compromised axonal cytoskeletal organisation is an early feature of FUS-associated ALS/FTD. We used an ALS-associated mutant FUS(P525L) and the FTD-mimic hypomethylated FUS, FUS(16R), to investigate the common and distinct cytoskeletal changes found in these two reported *Xenopus* models. Combining a novel atomic force microscopy (AFM)-based approach for *in vitro* cytoskeletal characterisation and *in vivo* axonal branching analysis, we found that mutant FUS reduced actin density in the dynamically remodelling growth cone, and reduced axonal branch complexity. We furthermore found evidence of an axon looping defect for FUS(P525L). Therefore, we show that compromised actin remodelling is potentially an important early event in FUS-associated pathogenesis.

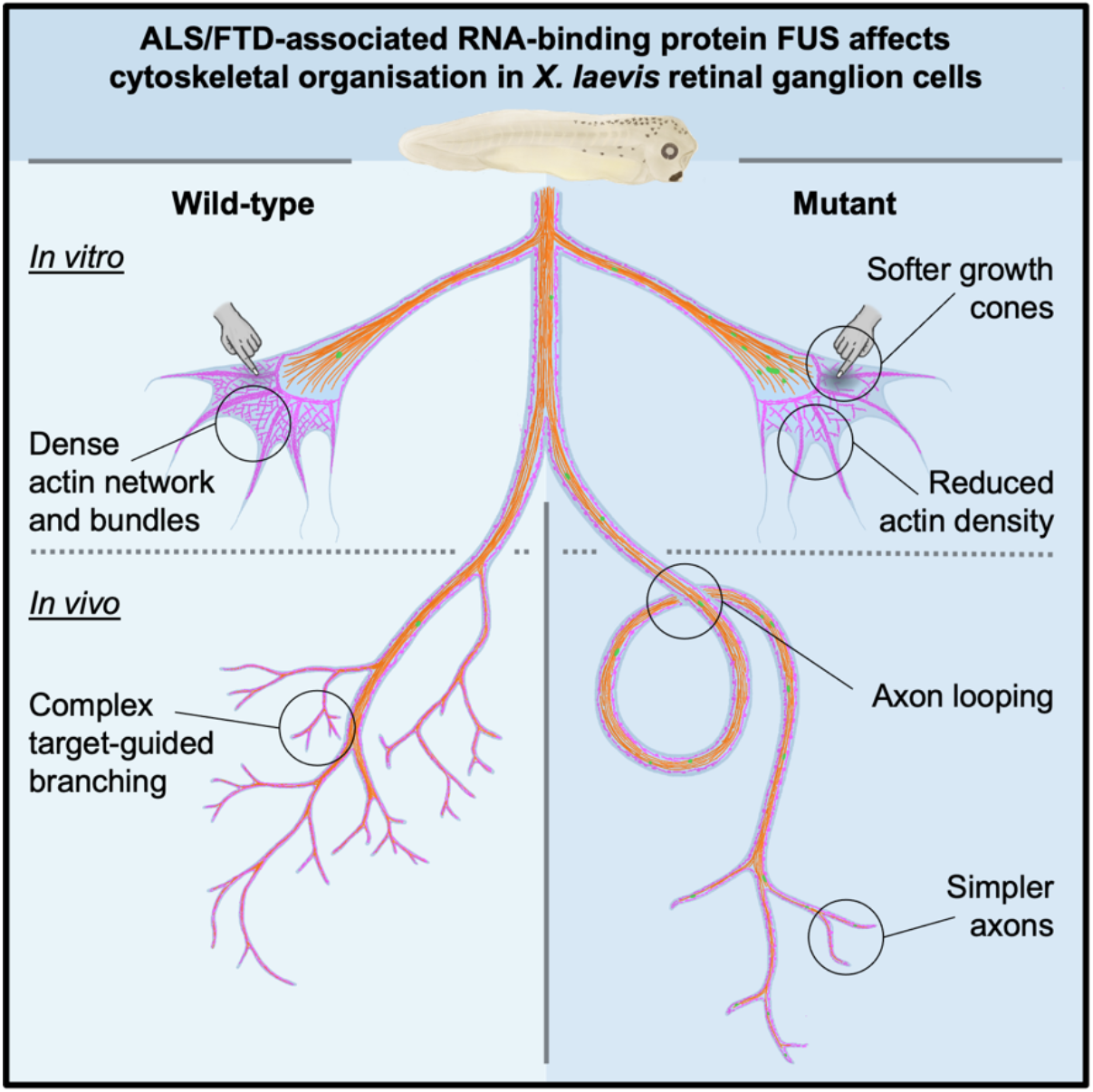

## Introduction

The RNA-binding protein (RBP) fused in sarcoma (FUS) forms cytoplasmic aggregates in variants of both amyotrophic lateral sclerosis (ALS) and frontotemporal dementia (FTD) (Kwiatkowski et al., 2009; Neumann et al., 2009; Vance et al., 2009). In some variants of familial ALS (fALS), mutation of FUS in the nuclear localisation sequence (NLS) leads to a rise in cytoplasmic FUS levels (Schoen et al., 2016). In FTD, wild-type FUS is instead found with far fewer methylated arginine residues (Dormann et al., 2012). This hypomethylation strengthens intramolecular cation-π interactions to accelerate irreversible FUS assembly formation (Hofweber et al., 2018; Qamar et al., 2018).

ALS/FTD has also been linked to changes in axonal cytoskeletal organisation. While ALS has been repeatedly linked to changes in axonal transport, less is known about changes in axonal actin, which are more challenging to study (Liu and Henty-Ridilla, 2022). Intriguingly, mutations in α-tubulin 4A are associated with familial ALS, ALS-FTD, and FTD (Mol et al., 2021; Smith et al., 2014), but patients with mutations in the actin polymerisation-regulating protein profilin-1 (PFN1) commonly display only ALS-like symptoms (Ingre et al., 2013; Wu et al., 2012). Therefore, actin and tubulin may be differently affected in different variants of ALS/FTD. Furthermore, cytoskeletal changes can affect FUS, providing potential mechanistic links between different familial ALS/FTD variants: PFN1 mutation causes disruption of nuclear envelope structure, likely via changed actin polymerisation, resulting in cytoplasmic mislocalisation of FUS (Giampetruzzi et al., 2019). In turn, FUS regulates dendritic actin by altering localisation of the mRNA encoding the actin-stabilizer protein NADH dehydrogenase subunit 1 (Nd1)-L (Fujii and Takumi, 2005). Inclusions of FUS also affect the detyrosinated glutamate microtubule network, compromising peripheral RNA localisation (Yasuda et al., 2017).

The relative contribution of actin and microtubule deregulation in ALS/FTD is under investigation (Liu and Henty-Ridilla, 2022), and has not been delineated for FUS-ALS/FTD. It has been argued that microtubules are particularly vulnerable in ALS, causing transport deficits (Clark et al., 2016; De Vos and Hafezparast, 2017), but also that the neuromuscular junction may be especially vulnerable to changes in actin, given that it is the largest synapse (Hensel and Claus, 2017). However, the coincident deregulation of actin and microtubules would have compounding effects on axonal health, as it would severely impact axonal structural remodelling. This would be apparent in axonal branching, which relies on both actin and tubulin rearrangement (Bodakuntla et al., 2021). This process is important for synaptogenesis (Kalil and Dent, 2014), and therefore of interest in understanding disease phenotypes The effects of NLS FUS mutants on unguided branching have been studied *in vitro*, with different results: branching was decreased in primary cortical cell axons (Groen et al., 2013) and motor neuron and cortical neuron dendrites (Qiu et al., 2014), but increased in hiPSC-derived motor neurons (Akiyama et al., 2019; Garone et al., 2021). As these studies focussed on identifying altered intracellular signalling pathways regulating branching, little is still known as to which cytoskeletal changes mutant FUS causes in developing axons. Furthermore, it is not known how FUS modulates axonal remodelling within a three-dimensional tissue environment, which contains chemical and mechanical cues that are critical for branch initiation and remodelling (Kalil and Dent, 2014).

In this study, we therefore tested the following hypothesis: altered cytoskeletal organisation is an early phenotype of FUS-associated ALS/FTD, and involves coincident deregulation of microtubules and actin filaments, resulting in compromised axonal branching. We use two previously reported ALS/FTD *Xenopus laevis* models: ALS-associated NLS mutant FUS(P525L) and the artificial FTD-mimic of hypomethylated FUS, FUS(16R), which contains sixteen strategically inserted arginine residues to increase the protein’s overall hypomethylation state (Qamar et al., 2018). We find that mutant FUS affects the actin cytoskeleton of the growth cone, the most dynamic axonal structure, without affecting axon shaft microtubules, resulting in selective softening of the growth cone. This is associated with compromised axonal branching complexity. We furthermore find evidence of an axon looping defect for FUS(P525L). Therefore, we show that compromised actin remodelling is potentially an important early event in FUS-associated pathogenesis.

## Results

### Mutant FUS consistently compromises growth cone but not axonal cytoskeletal integrity

To test for a perturbation in cytoskeletal organisation within the growth cone, we quantified the density of cytoskeletal filaments in distal axons expressing GFP, GFP-tagged FUS(WT), FUS(P525L), or FUS(16R) following mRNA injection (Supplementary Videos). To this purpose, we performed quantitative fluorescence microscopy on cultured RGCs by labelling filamentous actin (F-actin) with phalloidin, and β-tubulin by immunolabelling (Figure 1a). Microtubules were present in the central region of the growth cone and actin bundles and networks most strongly in the peripheral region (Figure 1b-c), as expected for neuronal growth cones (Omotade et al., 2017).

**Figure 1.**
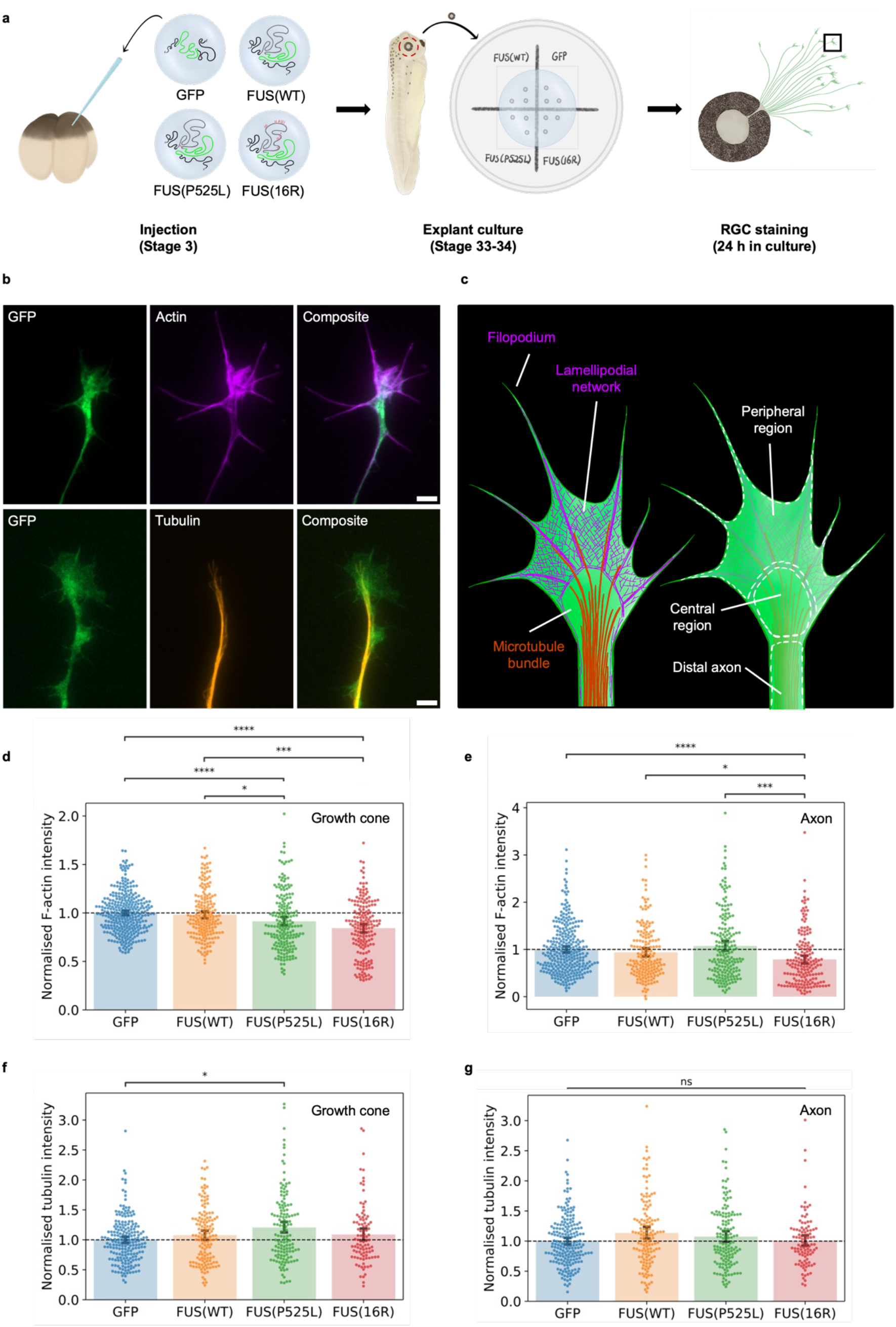
Mutant FUS expression affects actin density in distal axonal compartments. (a) Schematic of experimental procedure. mRNA encoding only GFP (green) or FUS (grey) fused to GFP with appropriate UTRs (black) is injected at the four-cell stage. Eye primordia are cultured at stage 33-34, and RGCs fixed and stained after overnight outgrowth. b) Sample images of growth cones labelled with phalloidin (F-actin) or an anti-β-tubulin antibody (tubulin)(scale bar: 5 μm). c) Schematic of growth cone cytoskeletal organisation. d) Normalised actin density in growth cone. e) Normalised distal axon actin density. f) Normalised growth cone tubulin density. g) Normalised distal axon tubulin density. (d-g): N >= 3 replicates for each condition, Kruskal-Wallis tests with Bonferroni correction.

Both FUS mutants caused changes in axonal actin density. Growth cone F-actin density was significantly reduced upon expression of FUS(P525L) and FUS(16R) compared with wild-type (WT) FUS or GFP-only controls (Figure 1d). Notably, this reduction was similar for both mutants. As growth cone area was unaltered (Supplementary Figure 1), this density change represents a decrease in total growth cone F-actin rather than a morphological change. In growth cones, actin remodelling and turnover are very rapid to facilitate directional migration (Omotade et al., 2017), so we wanted to know whether this fast remodelling was specifically affected. Therefore, we performed the same analysis on the distal axon shaft. We found that the reduction in F-actin caused by FUS(P525L) was confined to the growth cone, while FUS(16R)-induced F-actin disruption was also observed in axon shafts (Figure 1e).

Unlike for F-actin, tubulin density was not reduced by mutant FUS expression. In contrast, growth cone tubulin density was increased by FUS(P525L), relative to GFP only, but not by FUS(16R) (Figure 1f). Axonal tubulin density was unaltered for both FUS(P525L) and FUS(16R) (Figure 1g). Consistently with this, total RNA transport along axons was unaltered (Supplementary Figure 2), indicating axonal microtubule function was not compromised. Therefore, FUS(P525L) and FUS(16R) have overlapping but distinct effects on the axonal cytoskeleton, and affect the actin cytoskeleton more strongly.

### Cytoskeletal defects in mutant FUS-expressing growth cones alter mechanical properties

We next asked whether the observed changes in actin density compromised the mechanical properties of the growth cone. It is known that the dense actin network of the growth cone contributes considerably to the local stiffness (Xiong et al., 2009), while the stiffness of axon shafts is dominated by the microtubules (Ouyang et al., 2013). We therefore hypothesised that mutant FUS would alter the mechanical properties of the growth cone, but not those of axon shafts, based on our staining observations.

To test this hypothesis, we performed atomic force microscopy (AFM) on live growth cones and distal axon segments of RGCs *in vitro* (Figure 2a-b). On very thin and soft samples, such as growth cones, the properties of the material to which the sample is attached can considerably affect measurements (a phenomenon known as ‘substrate effects’). We therefore restricted our measurements to the thicker parts of the growth cone (commonly around 300 nm, cut-off of >150 nm), and refer to an ‘apparent Young’s modulus as determined by fitting a linearized Hertz model. We additionally validated that our approach could detect changes in mechanical properties upon changes in actin organisation, by acute treatment with cytochalasin D. This drug inhibits F-actin polymerisation and therefore disrupts the F-actin network. This treatment should result in a reduction in the apparent Young’s modulus of the growth cone, which we indeed observed (Supplementary Figure 3).

**Figure 2.**
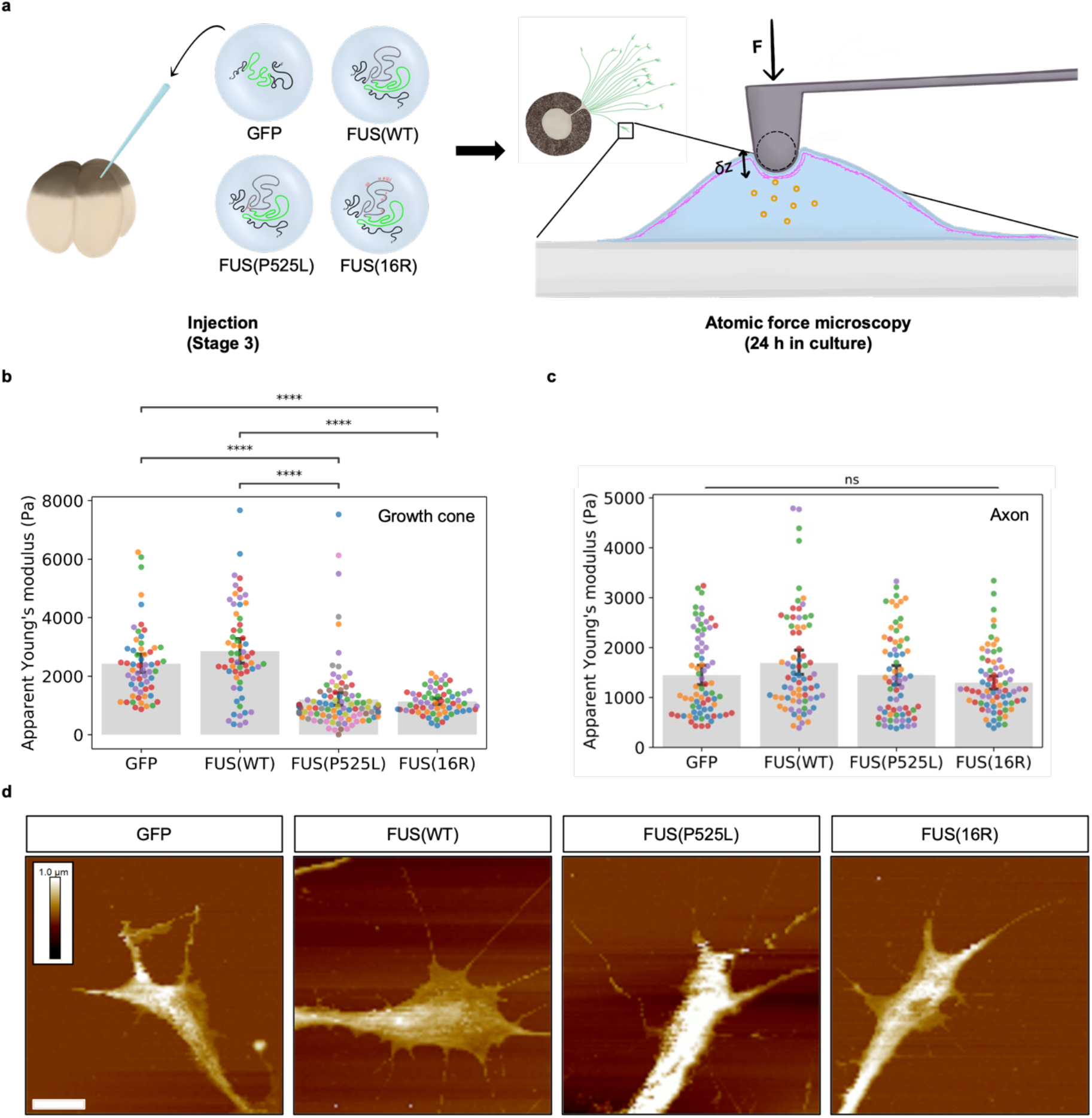
Mutant FUS reduces growth cone but not axon stiffness. a) Schematic of experimental procedure. RGCs are cultured as previously (Figure 1a). After outgrowth, force-displacement curves are measured for different parts of individual growth cones. b) Stiffness of growth cones expressing different FUS variants. c) Stiffness of axons expressing different FUS variants. d) Sample height maps of growth cones. Scale is the same for all images and shown for GFP-only condition: height is colour-coded, white scale bar (bottom left) is 5 μm. b-c) Colour labelling of data points indicates which are derived from the same axon or growth cone. Kruskal-Wallis tests with Bonferroni correction.

As expected, both mutant FUS variants affected the apparent Young’s modulus of growth cones, but not axons (Figure 2b-c). This occurred without changes in growth cone height (Figure 2d), and so indicates a change in cellular mechanical properties rather than a change in the contribution of substrate effects.

### Mutant FUS compromises axonal branching

We next investigated whether FUS’ effect on actin organisation was associated with functional defects later in development, by studying a key functional outcome of actin remodelling: branch formation at the target area. This process is critical for neuronal connectivity and is known to be compromised in a range of neurological disorders (Lin et al., 2021). *In vivo*, branch formation occurs in response to a complex set of target cell-derived cues (Bodakuntla et al., 2021), and so we studied this process *in vivo* to understand our findings’ physiological significance.

To investigate branch complexity, we quantified the morphology of RGC arbours *in vivo* at stage 45, when the axonal arbour has stabilised (Figure 3a). At this stage, dynamic cytoplasmic granules can be observed within the live embryo for FUS(P525L) and FUS(16R) (Supplementary Videos). We labelled individual axons by electroporation to allow arbour tracing.

**Figure 3.**
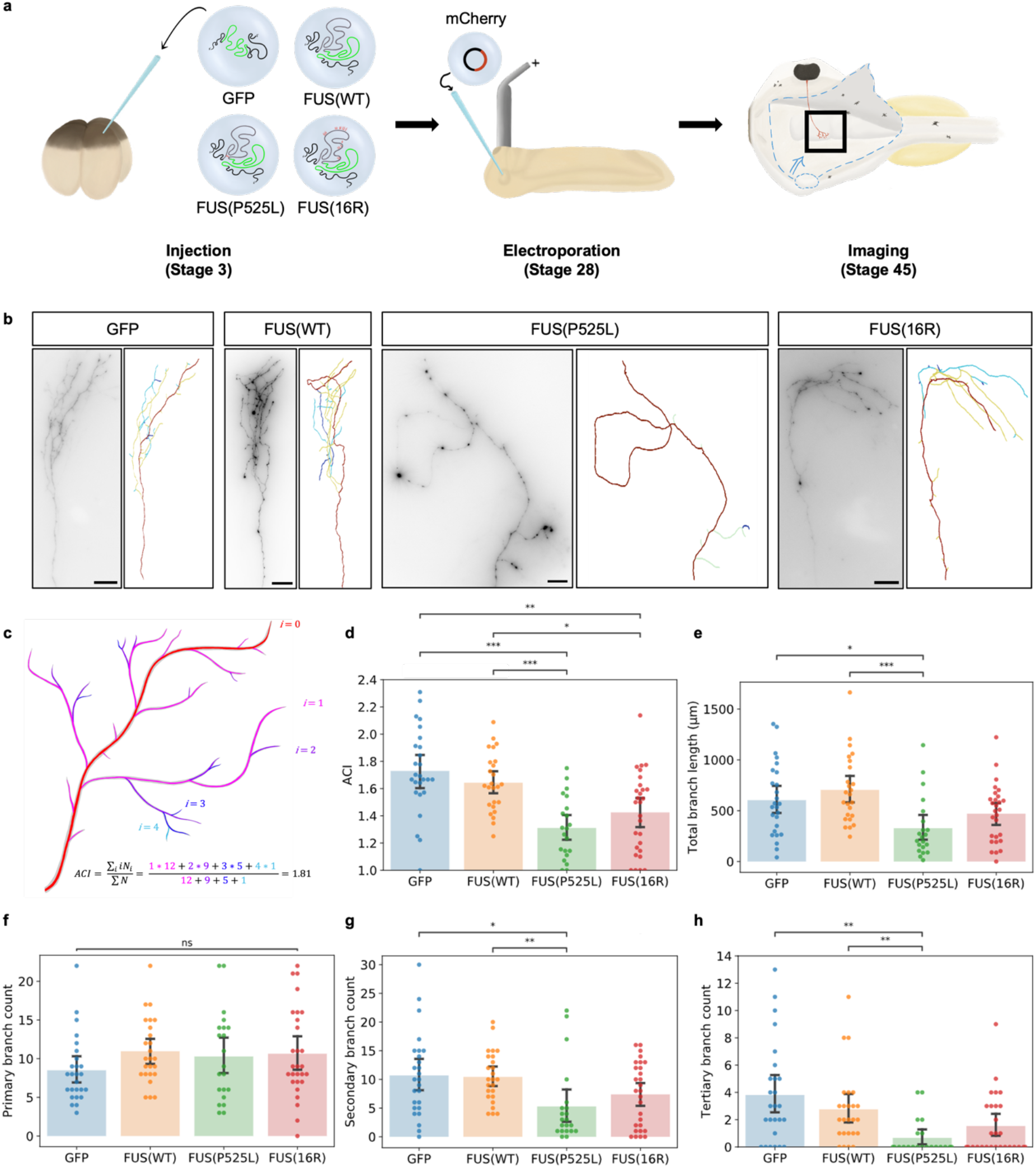
Axonal branching is compromised by mutant FUS expression. A) Schematic of experimental procedure. Embryos are injected as previously. At stage 28, the right eye primordium is electroporated with a plasmid encoding mCherry, resulting in sparse labelling. At stage 45, brains are exposed and RGC arbours in the tectum are imaged. B) Sample images of axons expressing different constructs. c) Schematic of calculation of axon complexity index (ACI). d) Axon complexity index. e) Total branch length. f-h) Number of primary, secondary, and tertiary branches per axon respectively.

Expression of both FUS(P525L) and FUS(16R) reduced axonal complexity compared with GFP-only and FUS(WT) controls (Figure 3b). Morphological complexity was quantified using the axon complexity index (ACI), a measure of the fraction of higher-order branches (Marshak et al., 2007)(Figure 3c). The average ACIs of stage 45 *X. laevis* RGCs have been reported to be in the order of 1.8 and 2.0, with an axon with ACI < 1.4 being considered simple (Cagnetta et al., 2019; Shigeoka et al., 2019; Wong et al., 2017). The ACI was significantly lower for axons expressing FUS(P525L) and FUS(16R), which were not significantly different from each other (Figure 3d). Overall, a reduction in branch length was observed for FUS(P525L)-expression axons compared with both GFP and FUS(WT)-GFP controls, but there was no clear effect for FUS(16R)-expressing axons (Figure 3e). The decrease in ACI associated with FUS(P525L)-expression was due to a decrease in the number of secondary and tertiary branches, as there was no reduction in primary branch numbers (Figure 3f-h).

### FUS(P525L) causes developmental transition defects

Notably, a subset of FUS(P525L)-expressing axons displayed an aberrant ‘looping’ phenotype (Figure 4a-b). While these axons reached the optic tectum without obvious errors in navigation, the main axon shaft executed an almost complete turn or loop within the tectum. We defined axons as ‘looping’ if they displayed this turning behaviour and were simple (ACI<1.4). This looping was rarely seen for axons expressing GFP, FUS(WT), or FUS(16R). The phenotype likely represents a failure of the axon to exit the elongation stage, i.e., a stop cue-dependent axon remodelling defect (Figure 4c). As tectal cells were also expressing mutant FUS, this phenotype may not be fully cell-autonomous. Therefore, while both FUS(P525L) and FUS(16R) compromise axonal arborisation, only FUS(P525L)-expressing axons display a developmental defect.

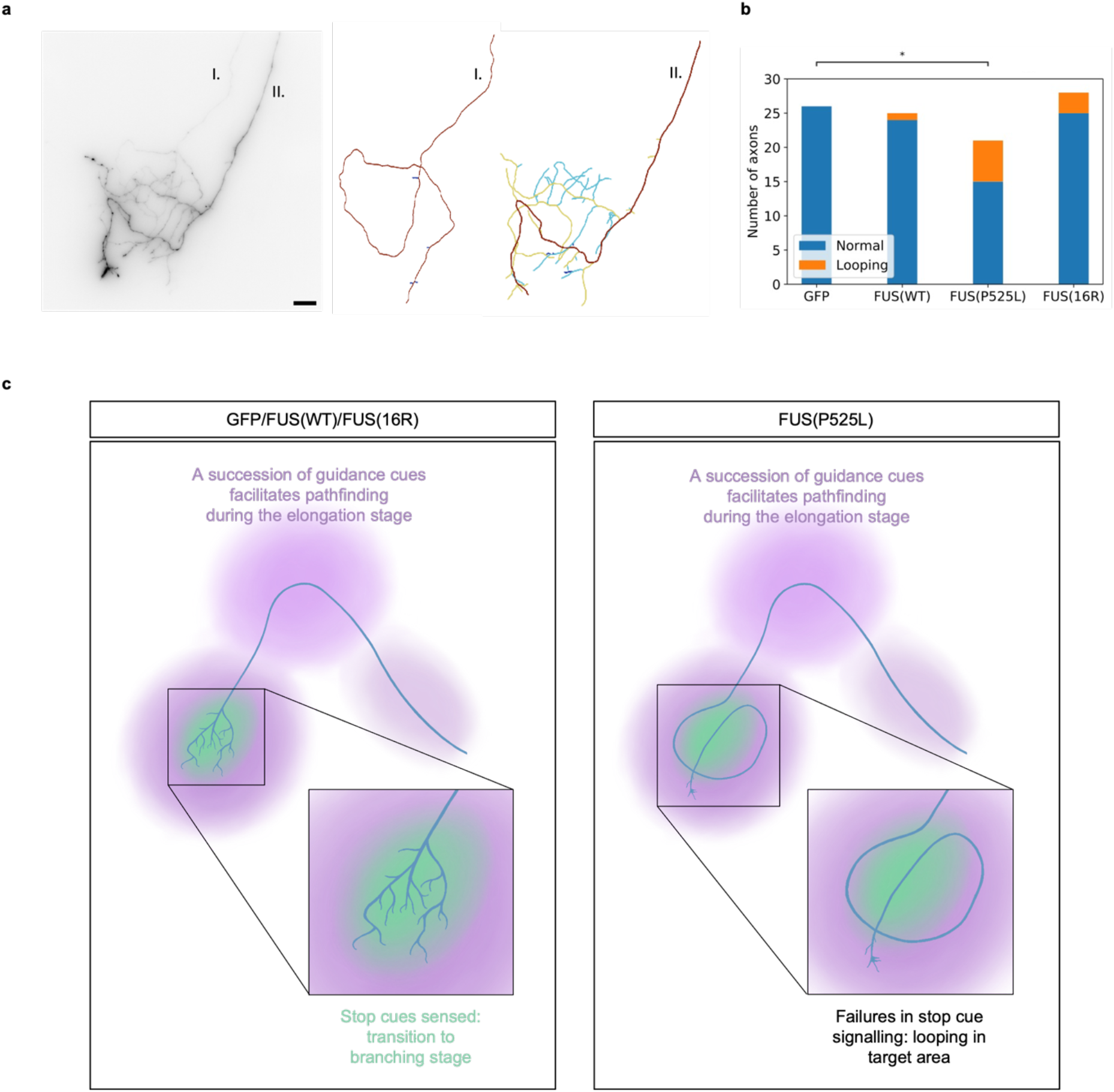
FUS(P525L) causes axon ‘looping’. A) Image of looping axon and normal axon within the brain of a FUS(P525L)-expressing embryo (scale bar: 20 μm). b) Number of looping axons. c) Schematic of looping defect as a developmental transition defect.

## Discussion

In summary, we observed changes in axonal actin cytoskeleton organisation upon expression of FUS(P525L) and FUS(16R), both *in vitro* and *in vivo*. This altered actin organisation could be downstream of changes in axonal protein synthesis and signalling pathways, but would also cause defects in these. Therefore, like altered axonal protein synthesis (Lin et al., 2021), altered actin remodelling could be a phenotype on which multiple mutations associated with ALS/FTD converge.

### FUS-induced actin changes may compromise growth cone mechanoproperties

We have shown that a combination of fluorescence imaging and AFM mechanical measurements permits us to quantify novel phenotypes associated with disease-linked FUS mutants. Given the complex, non-linear relationship between cellular mechanical properties and cytoskeletal organisation, our methodology is a promising method to assay the overall cytoskeletal integrity of the distal axon. However, technical limitations mean our AFM measurements represent changes in cellular mechanical properties, rather than direct measurements of cell stiffness (Young’s modulus).

Determination of Young’s modulus based on force curves was done using a linearised Hertz model, which assumes (1) that the sample’s thickness is infinite and (2) that the sample’s response to deformation is in the linear elastic regime (Dimitriadis et al., 2002). If indentation is more than ten percent of sample height, the first assumption does not hold, and substrate effects contribute to the calculated apparent stiffness (Domke and Radmacher, 1998). This is the case for our measurements, but as height was comparable for growth cones expressing different FUS mutants, it does not prevent a comparative rather than absolute study of the effects of FUS on cellular mechanical properties. If narrow probes are used to apply force, deformation is not in the low-strain regime, and the second assumption does not hold (Dimitriadis et al., 2002). We used conical probes with spherical tips of radius 70 nm. Therefore, the response we observe does not solely reflect Young’s modulus of sample and substrate, but also non-linear sample responses to high local strain. Therefore, we conclude that mutant FUS affects cellular mechanical properties, but do not determine that these are entirely or largely changes in Young’s modulus.

The changes in actin density observed in the growth cone are consistent with a reduction of Young’s modulus, so we hypothesise that a loss of growth cone stiffness contributes to the changes in mechanical properties we observe. The stiffness of filamentous actin networks derives from filament resistance to bending and stretching, and therefore filament density, length, and degree of crosslinking (Pegoraro et al., 2017). *In vitro* and computational studies show that Young’s modulus scales with filament density to a power of approximately 2.5 in cross-linked actin networks (Chen et al., 2020; Gardel et al., 2004; Pegoraro et al., 2017; Wang et al., 2020), and a correlation between actin density and Young’s modulus has been observed *in vivo* for the lamellipodia of fish keratocytes (Laurent et al., 2005). This hypothesis is further supported by parallel work showing depolymerisation of both actin and tubulin networks for FUS(P525L)-expressing mammalian cells, which are thicker samples, and this was correlated with a similar reduction in apparent Young’s modulus (Chung et al., 2022).

Changes in growth cone stiffness would be functionally significant and compromise development. These changes are likely functionally significant: growth cone stiffness is thought to be important for the transmission of tension from the peripheral to the central region (Xiong et al., 2009), which is important for growth cone advancement and steering (Suter et al., 1998). Furthermore, as filopodia often grow out of the lamellipodial network, they would produce a high local load on the actin network *in vivo*, as the extracellular matrix acts in a confining manner, making the Young’s modulus important for filopodial function (Chen et al., 2020). Furthermore, changes in the stiffness of nervous tissue overall could have non-cell-autonomous effects on axonal development: in *Xenopus*, the tectum where RGC axons branch is softer than tissues through which the axons navigate, and this facilitates axon unbundling, likely facilitating branching and synaptogenesis (Koser et al., 2016).

### FUS-associated cytoskeletal defects may arise through reduced axonal protein synthesis

A potential explanation for the defects in actin organisation may be FUS-induced reduction of axonal local protein synthesis (LPS). LPS and cytoskeletal reorganisation are co-dependent and co-regulated processes in axons, with cytoskeleton-associated proteins being locally synthesised and filaments acting as platforms for protein synthesis and regulation (Triantopoulou and Vidaki, 2022). For instance, it is known that local synthesis of β-actin at axonal branch points is required for complex arborisation (Wong et al., 2017). Several studies indicate that mutation of the FUS NLS inhibits global and local protein synthesis in neurons, including FUS(P525L) (Birsa et al., 2021; López-Erauskin et al., 2018; Murakami et al., 2015), and FUS(16R) is known to reduce axonal protein synthesis in *Xenopus* RGCs (Qamar et al., 2018). We observed that this reduction is not associated with changes in axonal RNA transport. Therefore, these changes in LPS are more likely due to changes in localisation of specific mRNAs or translational control, which could be due to altered FUS granule behaviour. In turn, this reduced LPS could partially explain the observed cytoskeletal defects, for instance if local synthesis of actin or actin-associated proteins is reduced.

*In vivo*, we observed defects in axon arborisation but not in axon pathfinding, which may be consistent with an impairment of LPS. RGC axonal branching has been demonstrated to be more sensitive to acute inhibition of local protein synthesis than axonal navigation (Wong et al., 2017): 30-minute acute inhibition of protein synthesis by treatment of exposed *X. laevis* brains with cycloheximide or anisomycin did not result in axonal navigation defects in somaless RGCs (stage 35-38), but did significantly reduce addition of filopodia and branches within the tectum (stage 41-43). Localised knockdown of *β-actin* mRNA translation in RGC axons resulted in a similar phenotype, indicating its translation within axons is important for this phenotype (Wong et al., 2017). Therefore, the branching phenotype observed here could be a downstream effect of reduced axonal LPS and associated remodelling of the axonal actin cytoskeleton. Higher-order branches may be more strongly affected due to differences in local energy or protein supplies, smaller branch diameters potentially affecting phase-separated structures restricting diffusion, or maturation defects in lower-order branches.

The observed defects in cytoskeletal organisation could in turn contribute to reduced LPS, especially *in vivo*, potentially explaining why mutations in cytoskeleton-associated proteins can result in similar ALS/FTD phenotypes as FUS mutations. Intact F-actin is known to be required for Netrin-induced LPS in *Xenopus* RGCs, with cytochalasin-D treatment preventing the early steps of translation initiation (Piper et al., 2015). Therefore, the FUS-associated reduction in actin density could directly compromise cue-dependent LPS *in vivo*, contributing to signalling defects.

### Altered signalling pathways may contribute to *in vivo* phenotypes

Signalling defects may contribute to FUS-induced loss of axonal complexity, which are captured by our *in vivo* approach to studying branching. In the *in vivo* experiments presented here, both tectal and retinal cells expressed mutant FUS, and so it remains possible that changes in tectal cells contribute to the observed phenotype. Extracellular stimuli have been shown to have major effects on axonal morphology, including guidance cues, growth factors, and morphogens (Cioni et al., 2018), and substrate stiffness has been reported to affect neurite branching *in vitro* (Flanagan et al., 2002). However, our *in vitro* data on cellular stiffness and morphology indicate growth cone cytoskeletal architecture is compromised upon mutant FUS expression, indicating the phenotype is at least partly cell-autonomous.

Our observations of a looping defect in FUS(P525L) also point towards a FUS-associated signalling defect compromising axonal cytoskeletal remodelling. In *X. laevis* RGCs, the growth cone halts advancement when it reaches the optic tectum, and does not overshoot its target area (Harris et al., 1987). Branching then occurs through the formation of filopodia at or near the base of the growth cone, notably also in axons that have been recently severed, and therefore independently of material from the soma (Harris et al., 1987). A defect in this pausing process could result in looping: the axon is able to sense guidance cues, and so is confined to the tectum upon arrival, but is unable to exit the advancement phase, resulting in looping within the defined volume of the tectum. The halt to extension of axons that reach their target area is thought to be mediated by “stop cues” (Cornel and Holt, 1992), and so signalling through these may be specifically altered (as opposed to signalling via guidance cues). Therefore, these data together suggest that while both FUS(16R) and FUS(P525L) affect the developing axon, FUS(P525L) may perturb specific LPS-based processes that guide axonal development. It has recently been shown that fast motor neurons expressing an ALS-associated variant of SOD1 are refractory to BDNF-mediated increases in axonal transport (Tosolini et al., 2022), which reinforces this idea, as BDNF also regulates axonal branching (Cohen-Cory et al., 2010). The role of these *in vivo* pathways and their changed regulation in ALS/FTD is an exciting new area of future study.

## Acknowledgements

This work was supported by UKRI Engineering and Physical Sciences Research Council (EPSRC) grants [EP/L015889/1] (to the Centre for Doctoral Training in Sensor Technologies and Applications, supporting F.W.v.T.) and [EP/H018301/1] (to C.F.K.), Wellcome Trust grants [3-3249/Z/16/Z] (to P.H.St.G.-H., C.F.K., and G.S.K.S.) and [089703/Z/09/Z] (to C.F.K.) and a Sir Henry Wellcome Postdoctoral Fellowship [215943/Z/19/Z] (to J.Q.L.) and a PhD studentship [109145/Z/15/Z] (to M.A.H.J.), the UK Medical Research Council (MRC) grants [MR/K015850/1] and [MR/K02292X/1] (both to C.F.K.), Infinitus (China) Ltd. (to C.F.K.), a European Research Council Consolidator award [772426] (to K.F.), a Michael J. Fox Foundation grant [16238], a Canadian Institutes of Health Research Foundation Grant and Canadian Consortium on Neurodegeneration in Aging Grant) [406915], a US Alzheimer Society Zenith Grant [ZEN-18-529769], the Alzheimer Society of Ontario Chair in Alzheimer’s Disease Research, and the UK Dementia Research Institute, which receives its funding from UK DRI Ltd, funded by the UK Medical Research Council, Alzheimer’s Society and Alzheimer’s Research UK.

This research was funded in part by the Wellcome Trust and Medical Research Council. For the purpose of open access, the authors have applied a CC BY public copyright license to any Author Accepted Manuscript version arising from this submission.

The authors would like to thank Asha Dwivedy, Yuri Efremov, and Meng Lu for assistance with experimental work.

## Author contributions

Conceptualisation, C.F.K., C.E.H., P.H.St.G.-H., F.W.v.T., J.Q.L.;

Methodology, F.W.v.T., L.C.S.W., I.M., J.Q.L., S.M., M.J., K.F., C.E.H., and C.F.K.;

Software, S.M. and M.A.H.J.;

Formal Analysis, F.W.v.T., L.C.S.W., I.M., and J.Q.L.;

Investigation, F.W.v.T., L.C.S.W., I.M., and J.Q.L.;

Resources, C.F.K., C.E.H., K.F., S.Q., and P.H.St.G.-H.;

Writing – Original Draft, F.W.v.T;

Writing – Review & Editing, C.F.K., J.Q.L., C.E.H., K.F., and L.C.S.W.;

Visualization, F.W.v.T., L.C.S.W., I.M., and J.Q.L.;

Supervision, C.F.K., C.E.H., J.Q.L., P.H.St.G.-H., G.S.K.S., and K.F.

## Methods

### Animal rights statement

This research has been regulated under the Animals (Scientific Procedures) Act 1986 Amendment Regulations 2012 following ethical review by the University of Cambridge Animal Welfare and Ethical Review Body (AWERB).

### Construct expression

To generate RGCs expressing different FUS variants of interest, conjugated to green fluorescent protein (GFP), *X. laevis* embryos were injected with *in vitro* synthesised mRNA. At the four-cell stage, blastomeres that will form the dorsal and ventral halves of the embryo are distinguishable, and injection of both dorsal blastomeres results in mRNA translation in both brain and eye tissue. Injected mRNAs encoded GFP fused to full-length wildtype FUS, FUS(P525L), or FUS(16R). A GFP-only control was also included. Injection was performed as described by Leung and Holt (2008). In brief, capped and polyadenylated mRNA was synthesised *in vitro* from plasmid stocks using the mMESSAGE mMACHINE™ SP6 transcription and poly(A) tailing kits (Invitrogen), which was subsequently diluted to a standard concentration of 200 ng/μL (100 ng/μL for GFP mRNA). Both dorsal cells of embryos at the four-cell stage were injected with 5 nL of mRNA solution.

### RGC culture

RGC culture was performed as described by Leung and Holt (2008). In brief, once embryos reached stage 33-34, eye primordia were dissected out and placed in No. 1.5 glass-bottom dishes (MatTek) that had been pre-coated overnight with poly-L-lysine (10 μg/mL, Merck) and subsequently for 1 hour with laminin (10 μg/mL in L15 medium, Merck). RGC cultures were kept in 60% L15 medium with 1X Antibiotic-Antimycotic (Gibco). Imaging was performed after overnight axon outgrowth.

### Quantitative fluorescence microscopy

RGC cultures were fixed with 2 % formaldehyde and 7.5 % sucrose in PBS for 20 min at RT. The samples were washed 5 times with PBS and permeabilized with 0.1 % Triton-X-100 in PBS for 5 min followed by 3 more washing steps with PBS. After blocking with 5 % donkey serum in PBS for 1 h, mouse anti-β-tubulin antibodies (ab131205, abcam) diluted 1:300 in blocking solution were applied and incubated overnight at 4°C. Dishes were washed with PBS 5 times for 3 min each before applying donkey anti-mouse Alexa Fluor 568 secondary antibodies (A10037, Invitrogen; 1:2000 diluted) and Alexa Fluor 647 phalloidin (A22287, invitrogen; 5 u/ml) in blocking solution for 1 h. Samples were washed again with PBS 5 times for 3 min each before imaging on a home-built widefield microscope. The microscope frame (IX83, Olympus) is equipped with a LED light source (DC4100, Thorlabs), a sCMOS camera (Zyla 4.2, Andor) and is controlled with the software Micro-Manager (Open Imaging). All images were acquired with a 60×/1.42 oil objective lens (PlanApoU, Olympus).

For quantitative fluorescence analysis, a custom written Matlab code was used to select individual growth cones. Only morphologically normal (non-collapsed) single growth cones were included in this analysis, to avoid selection bias. After background subtraction, intensity values were normalised to the signal area (i.e., the whole growth cone for actin staining, and the central region only for tubulin staining). Masks defining the area of interest were generated computationally following thresholding for actin, and by manual outlining for tubulin using Fiji (NIH).

### RNA granule tracking

To visualize RNA granule dynamics, RGCs were cultured and imaged on a spinning disk microscope. One-minute movies of axons were acquired (two frames per second, at constant exposure time and laser intensity). Kymographs were generated in Fiji (Schindelin et al., 2012), with a constant linewidth of 20 pixels. Granule paths were automatically detected and analysed using the KymoButler software (Jakobs et al., 2019). Distal ends of axons were imaged to enable imaging of unbundled and non-crossing axons, as well as unambiguous identification of anterograde and retrograde direction of motion in subsequent analysis (by presence of the growth cone).

### Atomic force microscopy

AFM measurements were performed using a commercial Bioscope Resolve atomic force microscope (Bruker, Santa Barbara). For the mechanical measurements, pre-calibrated Live Cell s (PFQNM-LC, Bruker AFM probes) cantilevers were used, with a nominal spring constant of 0.07 N/m s and the deflection sensitivity was measured at the start of each experiment using no touch calibration. The nominal radius of the probe is 70 nm. The microscope was operated in PeakForce QNM mode, data sampling rate was 0.25 kHz in most experiments and the force setpoint was selected so that the maximal indentation depth was around 200 nm. At least five cells per condition were measured and at least 10 force curves per cell region per condition were analysed. The extension part of the force curves (5 to 20% of the indentation) was fitted to a linearized Hertz model using Nanoscope 9.1 (Bruker), from which the Young’s modulus corresponding to each force curve was calculated. For pharmacological perturbation of the actin cytoskeleton, cultures were treated with 10 μg/mL of Cytochalasin-D for 30 minutes prior to measurement.

### Branching experiments

For branching experiments, injected embryos were electroporated at stage 26 (Wong and Holt, 2018). Electroporation was performed with a plasmid encoding mCherry (pCS2+ backbone). Two drops of solution (1 μg/μL) were delivered to the right eye primordium, quickly (<3s) followed by two pulses of 18 V (pulse width 50 ms, interval 1000 ms). This sparsely introduces the plasmid into retinal ganglion cells, resulting in a small number of labelled axons. Branching axons in the optic tectum were imaged using a 60X oil objective on a custom-built widefield microscope, as *z*-stacks with a spacing of 0.5 μm. To this purpose, the left (unelectroporated) eye and skin on the brain were removed, and embryos were mounted on permanox slides (Thermo Scientific) in frame-seal incubation chambers (Bio-Rad). All axons that were wholly and unambiguously traceable were included in the analysis (where up to three were labelled, or one or two axons were much brighter than other axons). As plasmid is occasionally electroporated into brain or spine cells, only axons within the correct orientation in the optic tectum were included in analyses. Analysis was performed using the SNT Neuroanatomy plugin for Fiji (Arshadi et al., 2021; Schindelin et al., 2012).

## Supplementary information

Supplementary videos for Figure 1: FUS localisation in RGC axons is granular. a) FUS(WT). b) FUS(P525L). c-d) FUS(16R). Live cultured RGC axons expressing GFP-tagged FUS constructs were imaged for one minute (one frame per second). Scale bar 5 μm, speed 5x.

**Supplementary Figure 1:**
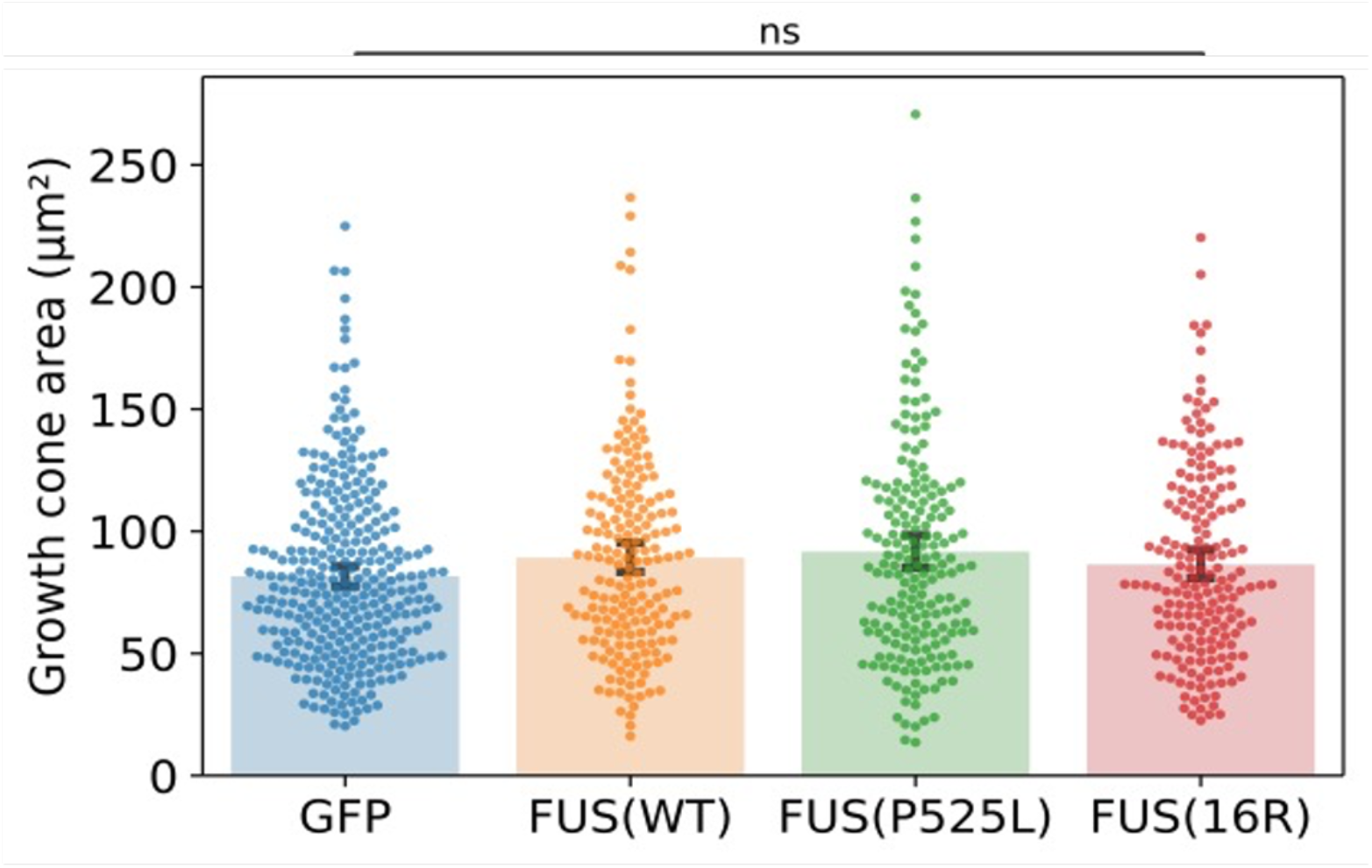
Growth cone area is not affected by expression of mutant FUS. Total area of phalloidin-stained growth cones (as in Figure 1b). N >= 3 replicates for each condition, Kruskal-Wallis tests with Bonferroni correction.

**Supplementary Figure 2:**
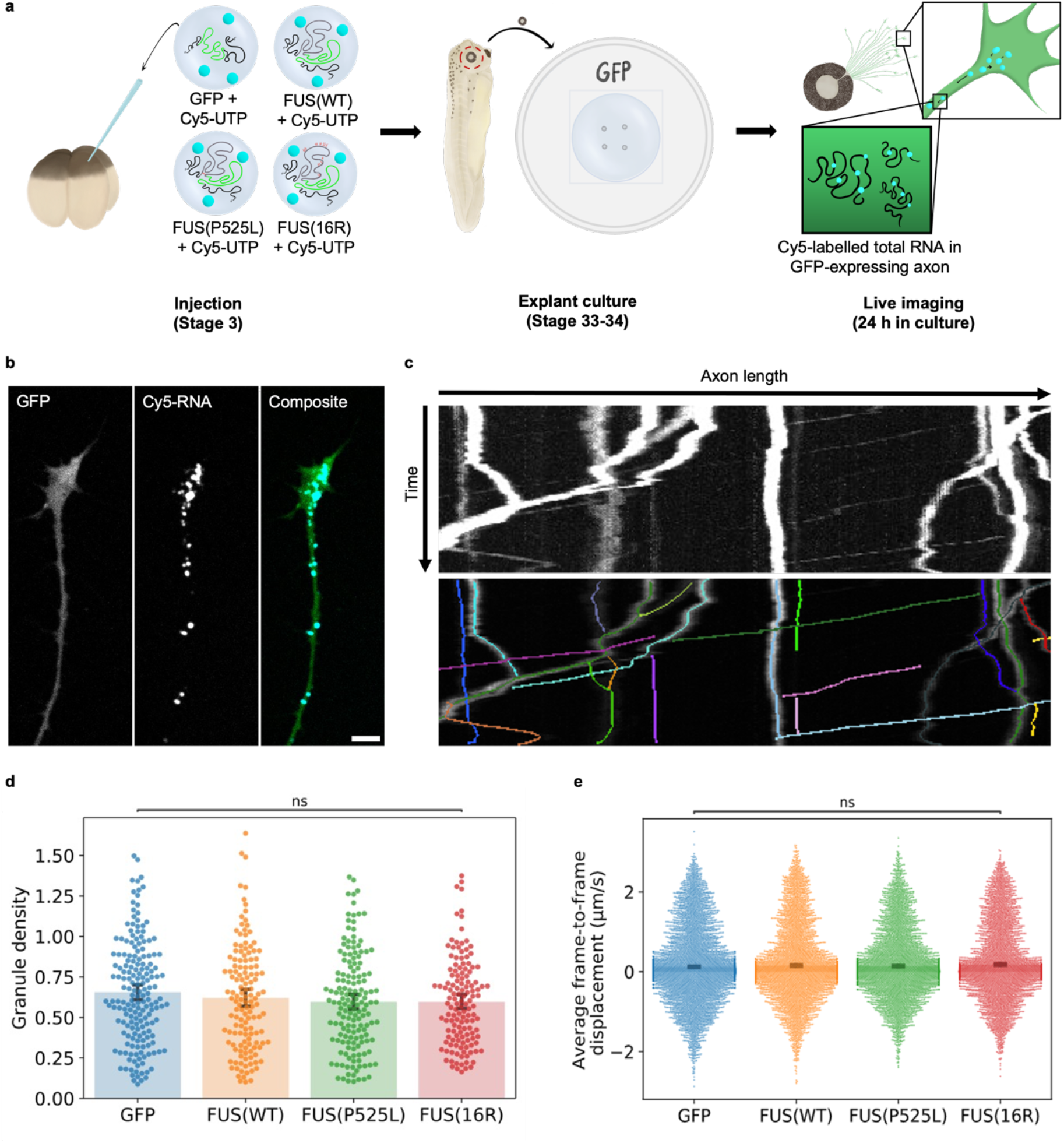
FUS expression does not alter axonal RNA transport dynamics. a) Sample preparation procedure. mRNA is co-injected with Cy5-UTP, which becomes incorporated into all embryonically synthesised RNA. RGCs are cultured as previously. Cy5-labelled RNA is visible as granules in the axon and growth cone (Piper et al., 2015; Wong et al., 2017). b) Sample image of RNA granules in GFP-expressing RGC axon (scale bar: 5 μm). c) Sample kymograph and track detection by KymoButler (Jakobs et al., 2019). d) Average granule density per unit length for individual axons. Data from three replicates. Total number of axons analysed: NGFP = 180; NFUS(WT) = 138; NFUS(P525L) = 155; NFUS(16R) = 136. E) Average frame-to-frame displacement for individual granules in 60 s. Data from three replicates. Total number of tracks identified: NGFP = 8587; NFUS(WT) = 6710; NFUS(P525L) = 7609; NFUS(16R) = 6202. Kruskal-Wallis tests with Bonferroni correction.

**Supplementary Figure 3:**
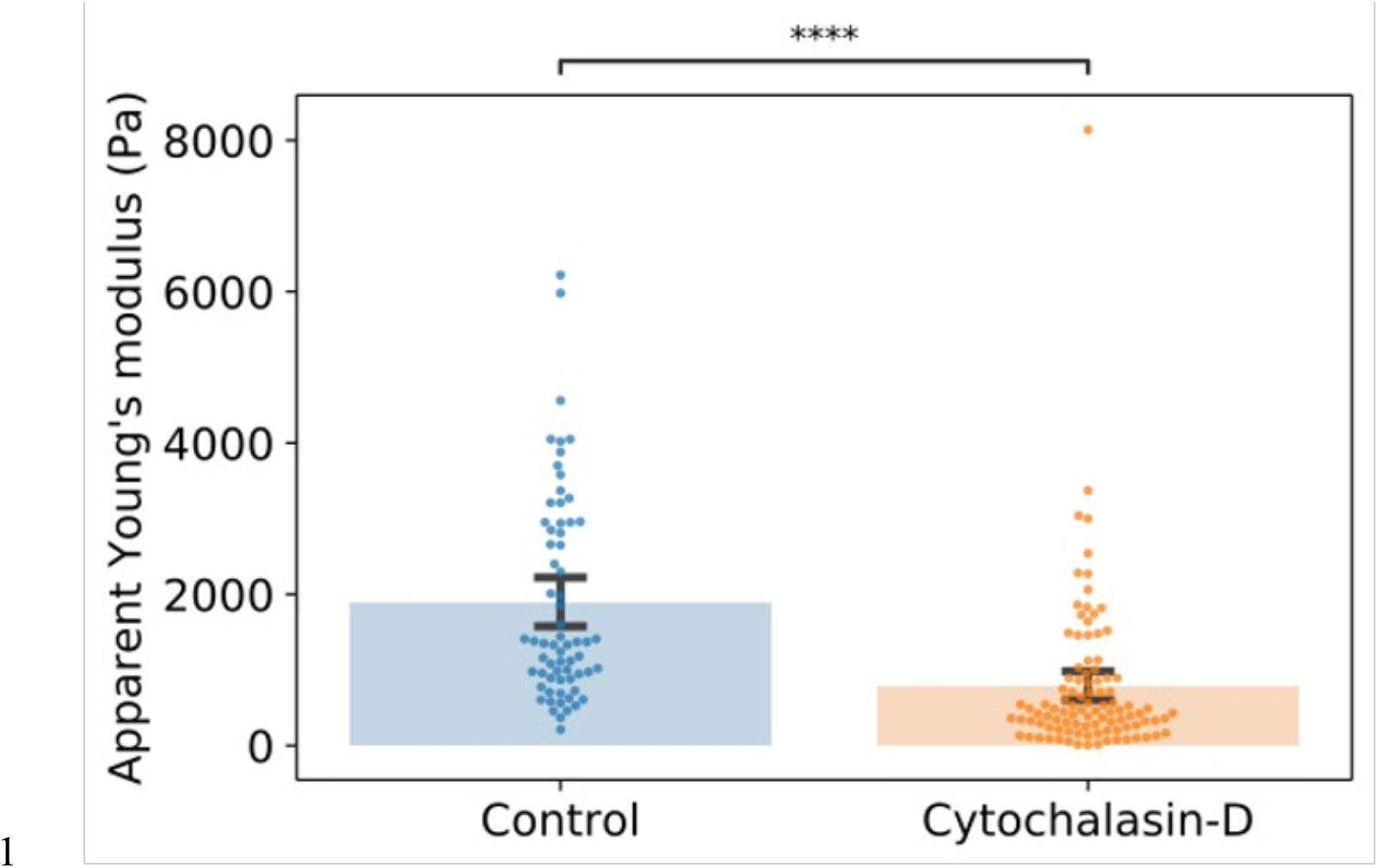
cytochalasin-D treatment lowers the apparent Young’s modulus.

Supplementary videos for Figure 3: Mutant FUS forms dynamic cytoplasmic granules in vivo. Z-stacks and single-plane videos of FUS-expressing cells were acquired towards the back of the brain and the top of the spine (away from the site of skin cutting). Expression was sparser in these areas than in the brain (as not all cells derive from dorsal blastomeres), allowing single cells to be readily distinguished. Scale bar 20 μm. a) FUS(WT)-GFP, brain; z = 1 μm. b) FUS(P525L)-GFP, back of brain; z = 0.5 μm. c) FUS(16R)-GFP, back of spine; z = 1 μm, d) FUS(P525L)-GFP, back of spine; Δt = 1 s, speed 5x. e) FUS(16R)-GFP, near top of spine; Δt = 1 s, speed 5x.

